# A Niclosamide-releasing hot-melt extruded catheter prevents *Staphylococcus aureus* experimental biomaterial-associated infection

**DOI:** 10.1101/2022.01.10.475592

**Authors:** Augusto Vazquez-Rodriguez, Bahaa Shaqour, Clara Guarch-Pérez, Emilia Choińska, Martijn Riool, Bart Verleije, Koen Beyers, Vivian J.A. Costantini, Wojciech Święszkowski, Sebastian A. J. Zaat, Paul Cos, Antonio Felici, Livia Ferrari

## Abstract

Biomaterial-associated infections are a major healthcare challenge as they are responsible for high disease burden in critically ill patients. In this study, we have developed drug-eluting antibacterial catheters to prevent catheter-related infections. Niclosamide (NIC), originally a well-studied antiparasitic drug, was incorporated into the polymeric matrix of thermoplastic polyurethane (TPU) via solvent casting, and catheters were fabricated using hot-melt extrusion technology. The mechanical and physicochemical properties of TPU polymers loaded with NIC were studied. NIC was released in a sustained manner from the catheters and exhibited antibacterial activity against *Staphylococcus aureus* and *Staphylococcus epidermidis* in different *in vitro* models. Moreover, the antibacterial efficacy of NIC-loaded catheters was validated in an *in vivo* biomaterial-associated infection mouse model using a methicillin-susceptible and methicillin-resistant strain of *S. aureus*. The released NIC from the produced catheters reduced bacterial colonization of the catheter as well as of the surrounding tissue. A sustained *in vivo* release of NIC from the catheters for at least 14 days was observed. In summary, the NIC-releasing hot-melt extruded catheters prevented implant colonization and reduced the bacterial colonization of peri-catheter tissue by methicillin sensitive as well as resistant *S. aureus* in a biomaterial-associated infection mouse model and has good prospects for preclinical development.

## 1. Introduction

Catheters represent an indispensable medical tool to improve the health quality and medical care of patients. The demand for catheters has increased due to the wide use of such devices in the administration of medications, nutritional support, blood sampling and performing dialysis in critically or chronically ill patients [1]. At least 150 million intravascular catheters are used annually in North America alone [2]. However, 3.5 % of these catheters are colonized by bacterial or fungal pathogens causing serious and costly bloodstream infections [3]. Catheter-related bloodstream infections (CRBSIs) increase the morbidity and mortality of intensive care patients with a mortality ranging from 19 to 34 % [4]. These infections are mainly caused by Gram-positive bacteria, mostly *Staphylococcus aureus* and *Staphylococcus epidermidis* [5, 6]. CRBSIs are primarily due to bacterial colonization of the catheter surface during insertion leading to a biofilm infection [5]. Bacterial cells encased in biofilm structures are very difficult to be eradicated by the immune defenses and antimicrobial agents [7, 8]. Hence, catheters must be removed and replaced to prevent further medical complications. Preventing bacterial attachment and subsequent biofilm formation on catheter surfaces would be the most cost-effective strategy to prevent CRBSIs [9].

There are many strategies described in the literature to address this increasing problem. These can be categorized as: 1) antifouling strategies such as hydration and steric repulsion, specific protein interactions, or low surface energy, and 2) antimicrobial mechanisms such as the release of biocidal agents, or surface microbicidal activity [10]. The release of biocidal agents approach has been extensively investigated over the past years via incorporating or coating medical devices with biocidal compounds such as antibiotics or other active compounds, for instance compounds identified in “repurposing” studies.

Antibacterial drug repurposing entails the use of already approved drugs for novel applications, such as antibacterial indications [11]. Niclosamide (NIC), an anthelmintic drug declared by the WHO as an essential medicine [12], has been identified as one of the most promising candidates to combat Gram-positive infections, especially those caused by *S. aureus* and *S. epidermidis*, being effective at relatively low concentrations (between 0.0625-0.5 μg/mL) [13–20]. However, given its poor solubility in water and low bioavailability, it may be less suitable for treating systemic infections [21–23]. Nevertheless, NIC has the potential to be applied to treat localized infections such as wound, soft tissue, gastrointestinal, skin and biomaterial associated infections (BAI) [14, 15, 23–27].

Based on this rationale, the aim of this research work was to develop an antibacterial catheter releasing NIC to prevent catheter-related infections. Specifically, we developed a local drug delivery system by incorporating NIC into the polymeric matrix of thermoplastic polyurethane (TPU) via solvent casting. Furthermore, the resulting material was used to fabricate antibacterial catheters by hot-melt extrusion (HME) technology. The produced catheters were characterized by studying the effect of NIC loading on the mechanical and thermal properties of the TPU. Moreover, the antimicrobial efficacy of the catheters was tested *in vitro* and subsequently in a murine BAI model.

## 2. Method and materials

### 2.1. Reagents

Thermoplastic polyurethane (TPU; Tecoflex EG-60D) was obtained from Lubrizol, VELOX, The Netherlands. The polymer contains a soft and hard segment in a ratio of 3:1. The glass transition temperature is 23°C [28]. Niclosamide (NIC; N3510, ≥98 % purity) and all other reagents and materials were purchased from Sigma-Aldrich (USA) unless indicated otherwise.

### 2.2. Bacterial strains

For *in vitro* microbiological evaluation, the bacterial strains used were the methicillin-susceptible *S. aureus* ATCC 25923 (MSSA), the methicillin-resistant *S. aureus* ATCC 33591 (MRSA), the methicillin-susceptible *S. epidermidis* O47 (MSSE) and the methicillin-resistant *S. epidermidis* ATCC 35984 (MRSE). The MSSA and MRSA were used in the *in vivo* BAI model. The bacteria were grown in tryptic soy broth (TSB; Difco Laboratories Inc, USA) at 37°C. Specific growth conditions for each experiment are described in the respective sections.

### 2.3. Material preparation

TPU was loaded with NIC at 2, 5 and 10 % (*w/w*) using the solvent casting approach adopted from the work of Shaqour *et al*. [29], as shown in *Figure 1*A. First, the required amount of NIC was suspended in chloroform and sonicated for 30 minutes. Then, a magnetic stirrer was used to homogeneously distribute the NIC particles for 30 minutes. Subsequently, the TPU was added to the system while stirring and the suspension was left overnight to dissolve the polymer. The ratio between TPU and chloroform was 12.5 % (*w/v*). The polymer solution with suspended NIC was then poured into a petri dish with a diameter of 200 mm to allow the chloroform to evaporate. Finally, the casted films were vacuum dried for 3 days at 25°C and 50 mbar.

**Figure 1.**
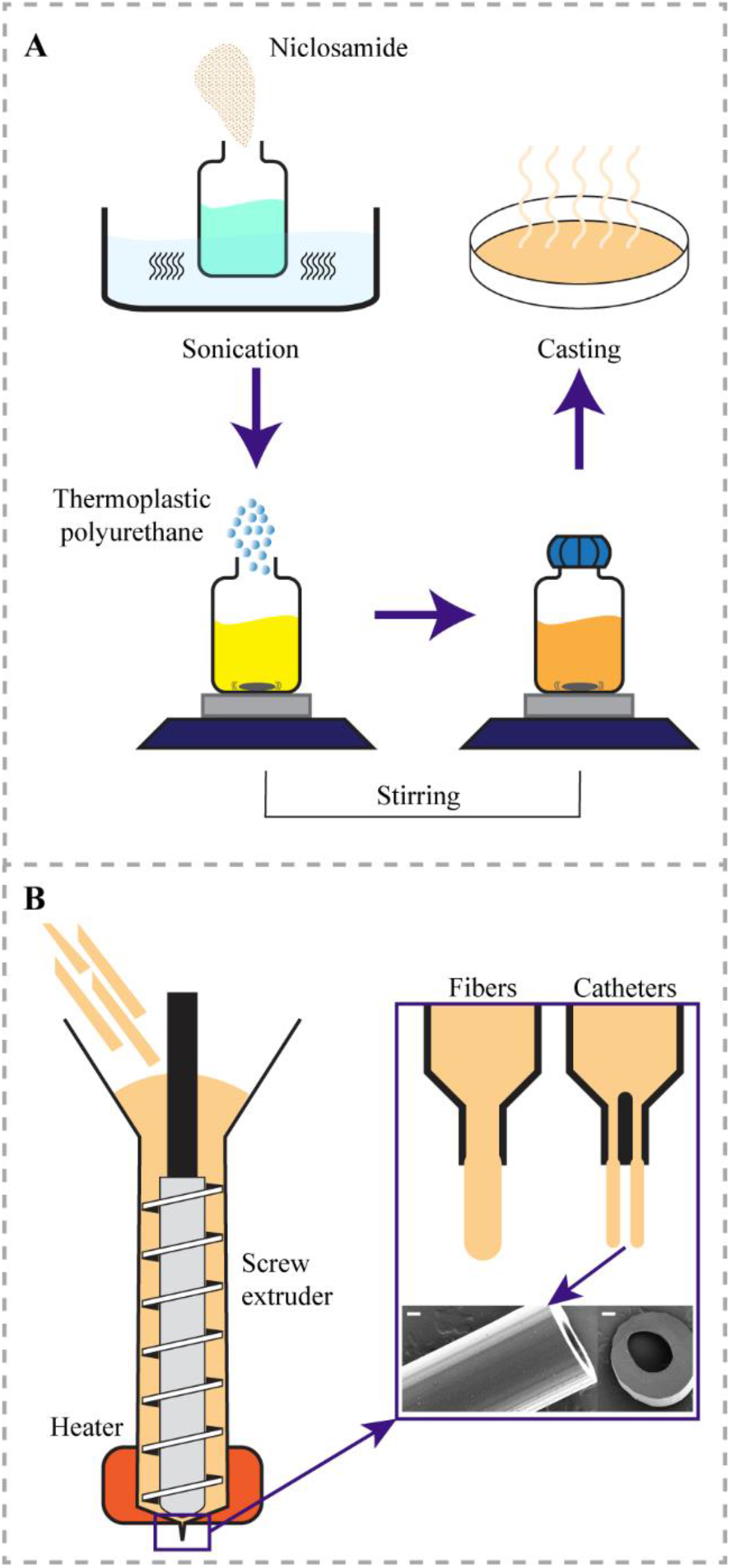
Process for A) incorporating NIC into the polymeric matrix of TPU using solvent casting and B) for producing fibers and catheters using HME technology. Scale bar in B is 200 μm.

Fibers and catheters were extruded using an in-house single screw extrusion setup attached with an e3d v6 stainless steel nozzle (E3D-Online, United Kingdom) with a diameter of 0.8 mm or a specially in-house designed and 3D-printed coaxial nozzle [30] (*Figure 1*B). To extrude the fibers and catheters, first, cast films (with or without NIC) were cut into stripes and directly fed into the extruder. Subsequently, fibers and catheters were extruded at a temperature of 180°C with a screw extrusion speed of 75 rpm. The extruded fibers were cut into 4 cm segments and used to evaluate the effect of NIC loading on the mechanical properties of the TPU, while the extruded catheters segments (0.5 and 1 cm length) were used for material characterization and for microbiological characterization. Catheter segments for microbiological testing were further sterilized by incubation in ethanol 70 % for 1 minute followed by air-drying for 30 seconds [29].

### 2.4. Microscopy

Catheter segments were inspected using an s9i microscope (Leica, Belgium). Catheters from each group were imaged and then analyzed using ImageJ software (v 1.52a; US National Institutes of Health, USA) by calculating the equivalent diameter (Equation 1) of the measured area for the inner and outer circles for each catheter (*n* = 6 for each group) [29].

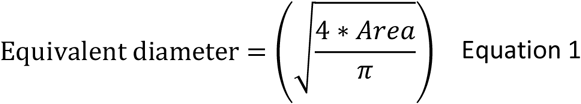

### 2.5. Mechanical testing

Tensile tests were performed according to ISO 527-1 standard [31] on an AGS 5 kND machine (Shimadzu, Germany). The tensile rate was set to 20 mm/min and the distance between grips was 20 mm. Extruded fibers (*n* = 5 per group) were used for this test. Then, the tensile stress at 100 % strain was measured.

### 2.6. Thermal analysis

Thermogravimetric analysis was conducted on a Q5000 analyzer (TA Instruments, USA). Samples of around 10 mg were placed on a platinum pan and then a dynamic heating ramp of 20°C/min with the resolution of 3°C to 500°C under nitrogen flow of 60 mL/min was applied.

### 2.7. Contact angle

The water contact angle measurements were done at room temperature using demineralized water and an OCA20 goniometer (Dataphysics, Germany). For each sample, the left and the right contact angles of at least 10 droplets with a volume of 2 μL were measured and averaged.

### 2.8. Fourier-transform infrared spectroscopy

NIC powder before and after heating to 180°C was examined using Fourier-transform infrared (FTIR) spectroscopy to study the effect of heat on NIC molecules. Moreover, to study the interactions between NIC and the TPU molecules, FTIR analysis was conducted on the produced catheters. The FTIR spectrometer used (Nicolet 8700, ThermoScientific, USA) was equipped with a diamond attenuated total reflectance (ATR) accessory. All ATR-FTIR spectra were recorded in the 400–4,000 cm^−1^ range at room temperature. The spectral resolution and accuracy were 4 cm^-1^ and ±1 cm^-1^, respectively.

### 2.9. X-ray Diffraction

X-ray diffraction (XRD) analysis was performed to analyze the solid-state characteristics of NIC pure powder, non-loaded TPU, and NIC-loaded TPU catheters. The XRD analyses were run in transmission mode on PANalytical X’Pert Pro X-Ray Diffractometer (PANalytical B.V., The Netherlands) equipped with an X’Celerator detector using a standard XRD method. The instrumental parameters used are listed in the supplementary Table 1.

### 2.10. *In vitro* drug release assay

Catheter segments loaded with NIC (2, 5 and 10 % (*w/w)*) and non-loaded, with a length of 1 cm were weighted using a microbalance (Sartorius, Germany). Then, each catheter segment was placed in 1 mL of phosphate buffered saline (PBS; ThermoFisher, USA) with 2 % of Tween 80 (ThermoFisher) and incubated at 37°C with an agitation of 120 rpm (*n* = 3 per group). The buffer solution was exchanged for the fresh solution at every time point (1, 3, 4, 6 and 24 hours, daily on days 2-10, and at 13, 16, 20 and 27 days). The aliquots were stored at −20°C for later use. The concentration of NIC released at every time point was calculated by measuring the absorbance at 340 nm of 300 μL of each aliquot in the flat bottom 96 well microtiter plates (Greiner Bio-One, USA) with a multi-well plate reader (Synergy H1, BioTek, USA). A calibration curve was plotted for NIC to estimate the concentration of drug released from the catheter segments. This calibration curve ranged from 1 to 50 μg/mL with R^2^ equal to 0.9998.

### 2.11. *In vitro* antimicrobial susceptibility test

To evaluate the minimal inhibitory concentration (MIC) and quantitatively assess the minimal bactericidal concentration (MBC) of NIC after heating at 180°C to mimic the extrusion process, MSSA was grown in TSB at 37°C and 120 rpm until reaching mid-logarithmic growth phase, and an inoculum of 1×10^6^ CFU/mL was prepared by dilution in fresh TSB, based on the optical density at 620 nm. Ninety microliters of TSB containing either unheated NIC control or NIC heated at 180°C were 2-fold serially diluted in TSB from 128 μg/mL to 0.125 μg/mL in a flat bottom microtiter plate. Immediately after, 10 μL of bacterial inoculum was added to the solution and incubated overnight at 37°C 120 rpm. As a control for bacterial growth, 10 μL of the inoculum was incubated in TSB without NIC. The MIC was defined as the lowest NIC concentration without visible bacterial growth overnight. For the MBC, 2 droplets of 10 μL of the undiluted samples from the wells without visible growth, were plated at blood agar plates (Oxoid, United Kingdom) and incubated at 37°C overnight. The MBC was assessed the next day as the lowest concentration of NIC which had caused ≥99.9 % reduction in numbers of CFU compared to the inoculum.

### 2.12. Evaluation of antimicrobial properties of TPU-catheters loaded with NIC

Catheter segments loaded with 2, 5, and 10 % (*w/w*) NIC and non-loaded catheter segments were analysed for antibacterial activity (release) and prevention of biofilm formation using the MSSA, MRSA, MSSE and MRSE strains.

Firstly, a modified Kirby-Bauer disk diffusion assay [32] was performed to determine the zone of inhibition (ZOI) of 0.5 cm catheter segments loaded with NIC and non-loaded catheters. Briefly, 1 to 3 colonies of each strain were incubated in 5 mL of TSB overnight at 37°C and 120 rpm. Two hundred microliters of an inoculum of 1×10^6^ CFU/mL was spread on blood agar plates with a cotton swab. The catheter segments were inserted vertically into the blood agar plates, which were then incubated at 37°C for 24 hours. The next day, the catheter segments were transferred to freshly inoculated blood agar plates and this step was repeated for 10 days. Each day the resulting zones of growth inhibition (including catheter diameter) were measured in mm.

Secondly, 1 cm catheter segments were placed in tubes containing suspensions of each of the 4 bacterial strains (approx. 5×10^6^ CFU/mL) in 1 mL of TSB. Catheter segments were incubated in the bacterial suspensions at 37°C and 120 rpm for 24 hours. After the incubation, 2 measures of bacterial growth were quantified: 1) planktonic bacterial growth in the medium; and 2) biofilm formation on the catheter surfaces.

To assess the antibacterial activity, planktonic cells in the medium were enumerated by quantitative culture, performed as follows: aliquots of the suspensions were taken, 10-fold serially diluted, and the dilutions and undiluted suspensions plated on tryptic soy agar (TSA) plates; these plates were incubated at 37°C for 24 hours and CFUs were determined. Similarly, biofilm formation was quantitatively measured by enumerating the viable cells attached to the surface. In brief, catheter segments were removed from the medium and rinsed 3 times in PBS to remove planktonic cells. Immediately thereafter, catheter segments were transferred to 1 mL of PBS, vortexed for 30 seconds, and sonicated for 15 minutes using an ultrasonic bath (Branson CPX2800-E, 40 kHz) to detach and disperse adherent-biofilm cells. This procedure does not affect viability of the bacteria. The bacteria recovered from the catheter segments were quantitatively cultured. For each experiment, triplicates were analyzed for each sample (NIC-loaded and non-loaded catheter segments).

Finally, scanning electron microscopy (SEM) was performed to visualize the morphological changes of MSSA bacterial cells adhered to NIC loaded and non-loaded catheter segments. As previously described in this section, bacterial suspensions were incubated with NIC loaded and non-loaded catheter segments until washing steps with PBS. Prior to SEM imaging, samples were fixed in a solution of 4 % (*v/v*) paraformaldehyde supplemented with 1 % (*v/v*) glutaraldehyde (Merck, USA) overnight at room temperature. Samples were rinsed twice with demineralized water for 10 minutes and dehydrated in a graded ethanol concentration series from 50 % to 100 % of ethanol. To reduce the sample surface tension, samples were immersed in hexamethyldisilane (Polysciences Inc., USA) overnight and air-dried. Before imaging, samples were mounted on aluminum SEM stubs and sputter-coated with a 4 nm platinum–palladium layer using a Leica EM ACE600 sputter coater (Microsystems, Germany). Images were acquired at 8 kV using a Zeiss Sigma 300 SEM (Zeiss, Germany) at the Electron Microscopy Center Amsterdam (ECMA; Amsterdam UMC, Amsterdam, The Netherlands). Of each tube, 6-8 fields were inspected and photographed at magnifications of 100× and 500×. Catheter segments loaded with 10 % of NIC were not included in the results as the material was partially degraded by the fixation procedure for SEM visualization.

### 2.13. *In vivo* evaluation of TPU-catheters loaded with NIC using a murine BAI model

The efficacy of NIC-loaded catheters to prevent infection was evaluated *in vivo* using a murine BAI model. As *S. aureus* is one of the most frequent pathogens in medical device-related infection [5, 6], the MSSA and MRSA strains were used for the *in vivo* evaluation of the NIC loaded catheters.

All experiments involving animals were carried out in accordance with both the European directive 2010/63/UE governing animal welfare and protection, which is acknowledged by the Italian Legislative Decree no 26/2014 and the company policy on the care and use of laboratory animals. All the studies were revised by the Animal Welfare Body and approved by Italian Ministry of Health (authorization n. 51/2014-PR). Male CD-1 mice (Charles River, Italy), weighing between 18-20 g, were housed in solid bottomed plastic cages with sawdust litter at a temperature of 20-22°C, a relative humidity range of 45-65 %, and lighting from approximately 06:00 to 18:00 hours daily. Mice were fed a standard maintenance diet (A. Rieper SpA, Italy) and drinking water was filtered from normal domestic supply. Diet regimen was ad libitum. A period of acclimatization of 5 days was implemented before any experimental procedure. Animals were monitored during the entire period of the studies and clinical signs were recorded.

Briefly, the mice were anesthetized using inhaled isoflurane at 2.5 % and the back of the mice was shaved using an electric razor and cleaned using an aqueous solution of benzalkonium chloride 0.175 % (*v/v*). Two small incisions were made in both flanks of the mice and two subcutaneous pockets were created using forceps, a 1 cm catheter segment was inserted into each pocket and the incisions were closed using surgical staples (AUTOCLIP™ wound closing system, Clay Adams, USA) under sterile conditions. Each mouse received two catheter segments with the same concentration of NIC, either non-loaded, 2 or 5 % (*w/w*) NIC loaded catheters (*n* = 7 mice per group, so *n* = 14 catheter segments per group). Three additional mice were added to each experimental group and dedicated to histological analysis on day 3 post-infection, similarly at the same time point, a group comprised of mice implanted with TPU catheters without infection was also added for comparison in the histological analysis (Figure 2).

**Figure 2.**
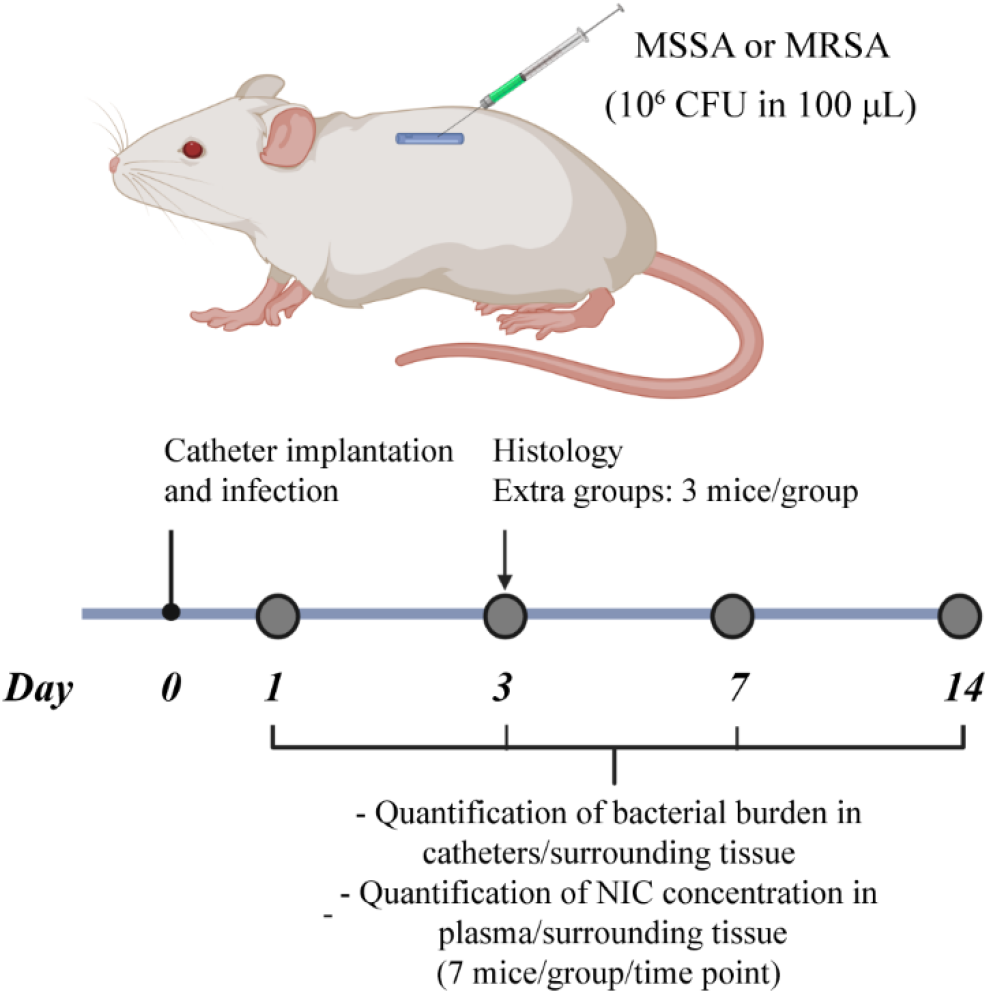
*In vivo* efficacy evaluation of non-loaded, 2 and 5 % (*w/w*) NIC loaded TPU catheters using a murine BAI model. Two 1 cm catheters segments were implanted subcutaneously in the mice and a bacterial challenge (10^6^ CFU in 100 μL) of either MSSA or MRSA was injected intraluminally into each catheter. Blood, catheter segments, and the surrounding tissue were collected of mice sacrificed on day 1, 3, 7 and 14 after challenge. Bacterial colonization of the catheter and the surrounding tissue, and the NIC concentration in plasma and in the tissue surrounding the catheter were quantified.

Immediately following catheter implantation, 100 μL containing 10^6^ CFU of either MSSA or MRSA were injected intraluminally into the catheter segments (**Figure 2**).

Animals were anesthetized using inhaled isoflurane (2.5 %) and the blood collected by terminal cardiac puncture at 1, 3, 7 or 14 days post-infection. Blood, catheter segments and the respective surrounding tissue were collected. Biopsies were aseptically retrieved using a 12 mm diameter biopsy punch and accordingly processed for bacterial load determination: catheter segments were separated from their surrounding tissue and rinsed3 times in 0.9 % saline solution to remove non-adherent bacteria. Next, catheter segments were transferred into tubes containing 1 mL of sterile PBS, vortexed for 30 seconds and sonicated for 15 minutes using an ultrasonic bath (Branson CPX2800-E, 40 kHz) to detach and disperse biofilm cells from the catheter surfaces. Recovered bacteria from the catheters were then quantitively cultured. Surrounding tissue was homogenized in 1 mL of PBS using a Precelys tissue homogenizer device (Bertin Technologies, France). Tissue homogenates were quantitatively cultured to enumerate viable bacterial cells residing in the tissue. In case of bacterial growth on day 14 post-infection in the experiment with MRSA, *S. aureus* colonies were retrieved from all of the experimental groups and the MIC for NIC was determined for all single retrieved colonies following the CLSI guidelines[33, 34].

For histopathology analysis, biopsies were retrieved from histology dedicated mice at day 3 post-infection and preserved in 10 % neutral buffered formalin. Following fixation, samples were embedded in paraffin, sectioned at a nominal thickness of 3-5 μm and stained with hematoxylin-eosin for histological analysis.

The NIC *in vivo* release profile in animals carrying NIC-loaded catheters was determined in plasma and in the tissue surrounding the catheter segments collected on day 1, 3, 7 and 14 post infection. First plasma was collected as follows: 1 mL of blood was placed in K3-EDTA collection tubes (Greiner Bio-One, USA) and centrifuged at 3,000 RCF, 4°C for 10 minutes. Fifty microliters of supernatant-containing plasma were then retrieved and mixed with 150 μL of HEPES 0.1 N and stored at −20°C until analysis.

Second, catheter surrounding tissue was processed as previously described and resulting tissue homogenates were further homogenized by enzymatical digestion: 1 mL of a solution of collagenase type 1 (8 mg/mL in PBS) and incubated for 3 hours at 37°C with agitation. Digested tissue homogenates were kept at −20°C until analysis. Prior to the analytical procedure, plasma and tissue homogenate samples were deproteinized with 2 volumes of acetonitrile containing diclofenac (200 ng/mL) as internal standard, vortexed and then centrifuged at 3,000 RPM for 10 minutes. After centrifugation, supernatants were diluted (160 μL ultrapure water + 200 μL of supernatant) using a Hamilton Microlab STARlet small liquid handler (Hamilton Company, Switzerland). Levels of NIC in supernatants were determined by analyzing 2 μL of the supernatants using a Waters ultrahigh performance liquid chromatography system (UPLC; Milford, USA) coupled with an API400 (Applied Biosystems/MDS Sciex, USA)) in tandem mass spectrometry mode (LC-MS/MS). Chromatographic separation was performed using a Waters Acquity UPLC BEH C18 30×2.1 mm, 1.7 μm analytical column. Mobile Phase A consisted of 0.1 % (*v/v*) formic acid in water; mobile phase B was 0.1 % (*v/v*) formic acid in acetonitrile.

### 2.14. Statistical analysis

Quantitative data was expressed as the average ± standard deviation, with the number of samples stated in each experiment. The statistical analysis of all the *in vitro* and *in vivo* characterization experiments was performed using a one-way analysis of variance (ANOVA) with Dunnett’s comparison test to evaluate post-hoc differences between the test groups with respect to the control group. Alpha was set at 0.05 for all analyses. Statistical analysis was performed using Prism8 (GraphPad Software, San Diego, CA, USA). * Indicates a *p*-value of 0.01 to 0.05, ** indicates a *p*-value of 0.001 to 0.01, *** *p*-value of 0.0001 to 0.001, **** indicates a *p*-value < 0.0001.

## 3. Results

### 3.1. TPU films loaded with NIC and catheter/fiber extrusion

The NIC loaded and non-loaded TPU films were successfully produced. The non-loaded TPU films were transparent while the NIC-loaded ones were yellowish and opaque. TPU is highly soluble in chloroform[21]. The level of coloring and opaqueness increased with increasing NIC content. However, the NIC distribution was homogeneous as no agglomerates were spotted in the films.

The stress at 100 % strain was used to assess the mechanical properties of the produced fibers and whether the addition of NIC affected TPU’s mechanical properties. The extruded fibers from the 0.8 mm nozzle had a diameter of 0.91 ± 0.08 mm. There were no significant differences between the NIC-loaded and non-loaded fibers and tubes in term of diameter dimensions. The generated stress-strain curves for TPU and drug loaded samples are shown in supplementary figure 1. The stress at 100 % strain was 11.1 ± 0.3, 10.7 ± 0.5, 11.4 ± 0.5 and 9.9 ± 0.4 MPa for the non-loaded and 2, 5 and 10 % (*w/w*) NIC loaded TPU fibers, respectively (*Figure 3*A). Only the 10 % (*w/w*) NIC loaded fibers showed a slight but significant decrease in the stress (*p* = 0.0012) when compared to non-loaded TPU fibers, which may affect the mechanical properties of extruded catheters.

**Figure 3.**
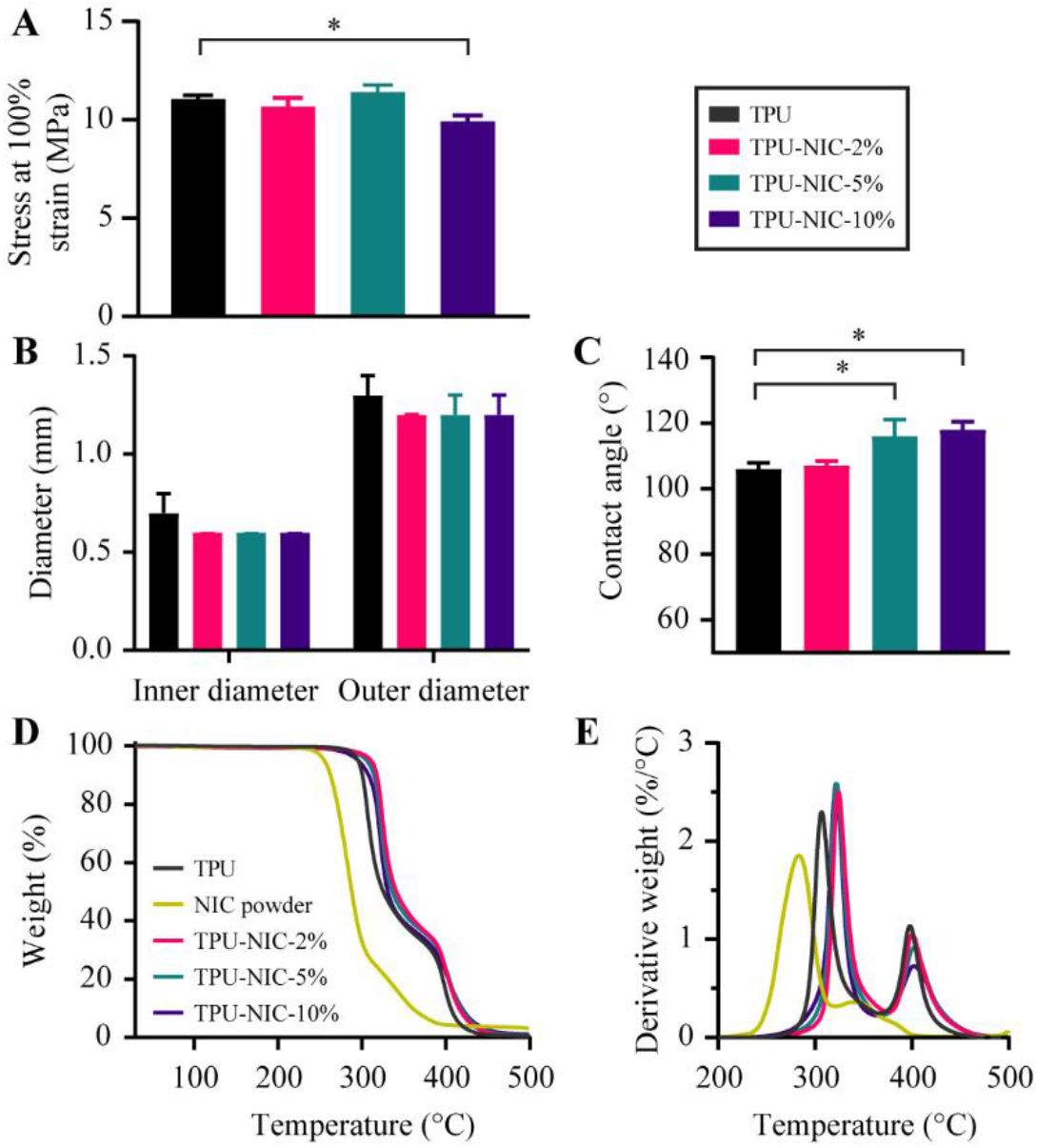
(A) Stress at 100 % strain for produced TPU and NIC loaded TPU fibers (*n* = 5), (B) Inner and outer diameter of produced TPU tubes (*n* = 6), (C) Water contact angle measurements of produced TPU catheters (*n* = 10). (D) Percentage weight loss vs. temperature and (E) Derivative of percentage weight loss vs. temperature in produced TPU fibers. * Indicates a *p*-value of 0.01 to 0.05.

The produced catheters had an average outer and inner diameter of 1.24 ± 0.62 mm and 0.62 ± 0.01 mm, respectively, with no differences between the different types of catheters (*Figure 3*B). The results of water contact angle measurements from non-loaded and NIC-loaded TPU catheters are shown in *Figure 3*C. All catheters were hydrophobic (*i.e*. contact angle higher than 90°) [35], with a contact angle of 106.5 ±1.9°, 106.8 ± 1.5°, 116.3 ± 5.0° and 118.4 ± 2.5° for non-loaded, 2, 5 and 10 % NIC loaded TPU catheters, respectively. The hydrophobicity of the NIC-loaded catheters increased with higher NIC loading, which is expected since NIC is considered a hydrophobic molecule [36]. However, only the 5 and 10 % (*w/w*) NIC-loaded catheters (*p* < 0.0001) showed a significant increase in the contact angle when compared to non-loaded catheters.

Thermogravimetric analysis (*Figure 3*D and E) showed the thermal degradation behavior for NIC powder, non-loaded and NIC-loaded TPU catheter segments. NIC and TPU had an onset degradation temperature (T_onset_) of 260°C and 295°C, respectively. Measurements of the NIC loaded catheters showed that the T_onset_ for NIC-loaded TPU was slightly higher than that of non-loaded catheters. T_onset_ for 2, 5 and 10 % NIC loaded TPU were 315°C, 313°C and 311°C, respectively. Such increase in T_onset_ was also reported when TPU was loaded with ciprofloxacin [29] or tetracycline hydrochloride [37], which indicates that, like NIC, the presence of these antibiotics improved the thermal stability of the polymer.

### 3.2. Fourier-transform infrared spectroscopy and X-ray Powder Diffraction analysis

The FTIR analysis for the untreated NIC powder and NIC heated at 180°C revealed no major changes in the spectra due to the heating (supplementary figure 2). The FTIR spectra of NIC powder, non-loaded and NIC loaded TPU are shown in Figure 4A. The NIC spectrum showed characteristic bands at 3,576.34 cm^-1^, 3,088.44 cm^-1^ and 1,679.69 cm^-1^ corresponding to -OH, -NH, and C=O groups, respectively (*Figure 4*B) [38]. The non-loaded TPU tubes exhibited a typical distribution of absorption bands for this type of material [39].The vibrational band at 3,330 cm^-1^ corresponds to NH stretching vibrations. The peaks in the range from 3,000 to 2,800 cm^-1^ relate to CH asymmetrical and symmetrical stretching vibrations [39]. The double peak in the region of 1,680 to 1,740 cm^-1^ (C=O stretching vibrations), the bands above 1,500 cm^-1^ (probably N-H bending vibrations and C-N stretching vibrations), and the strong bands in the region of 1,300 to 1,000 cm^-1^ (asymmetrical and symmetrical O-C-O stretching vibrations) are associated with the bonds of urethane groups [39]. The FTIR spectrum of NIC-loaded TPU tubes has characteristic peaks similar to those of the spectrum of non-loaded TPU tubes (*Figure 4*A). Moreover, also the new specific peaks due to the presence of NIC appeared in the spectra of the loaded with 5 % and 10 % (*w/w*) NIC. These peaks characteristic for NIC are practically invisible at the spectrum with the lowest NIC content. It was probably caused by the low intensity of these bands and the fact, that they were overlapped by the TPU peaks. For the higher content of NIC, the intensity of a peak of hydrogen-bonded N-H group in the amorphous region of TPU (3,320 cm^-1^) significantly decreased and the band of free N-H group with no H-bond formation observed at 3,450 cm^-1^ became stronger. Probably the drug particles weakened the force between H-bond groups of TPU and hindered the formation of a new H-bond, a similar effect was observed for the thermally processed composite of montmorillonite and polyamide [40]. Additionally, the strong band at 3,535 cm^-1^ was observed, which can be assigned to the hydroxyl groups of NIC. In the comparison to the pure drug, the intensity of this peak increased, due to the amorphization of the drug[41].

**Figure 4.**
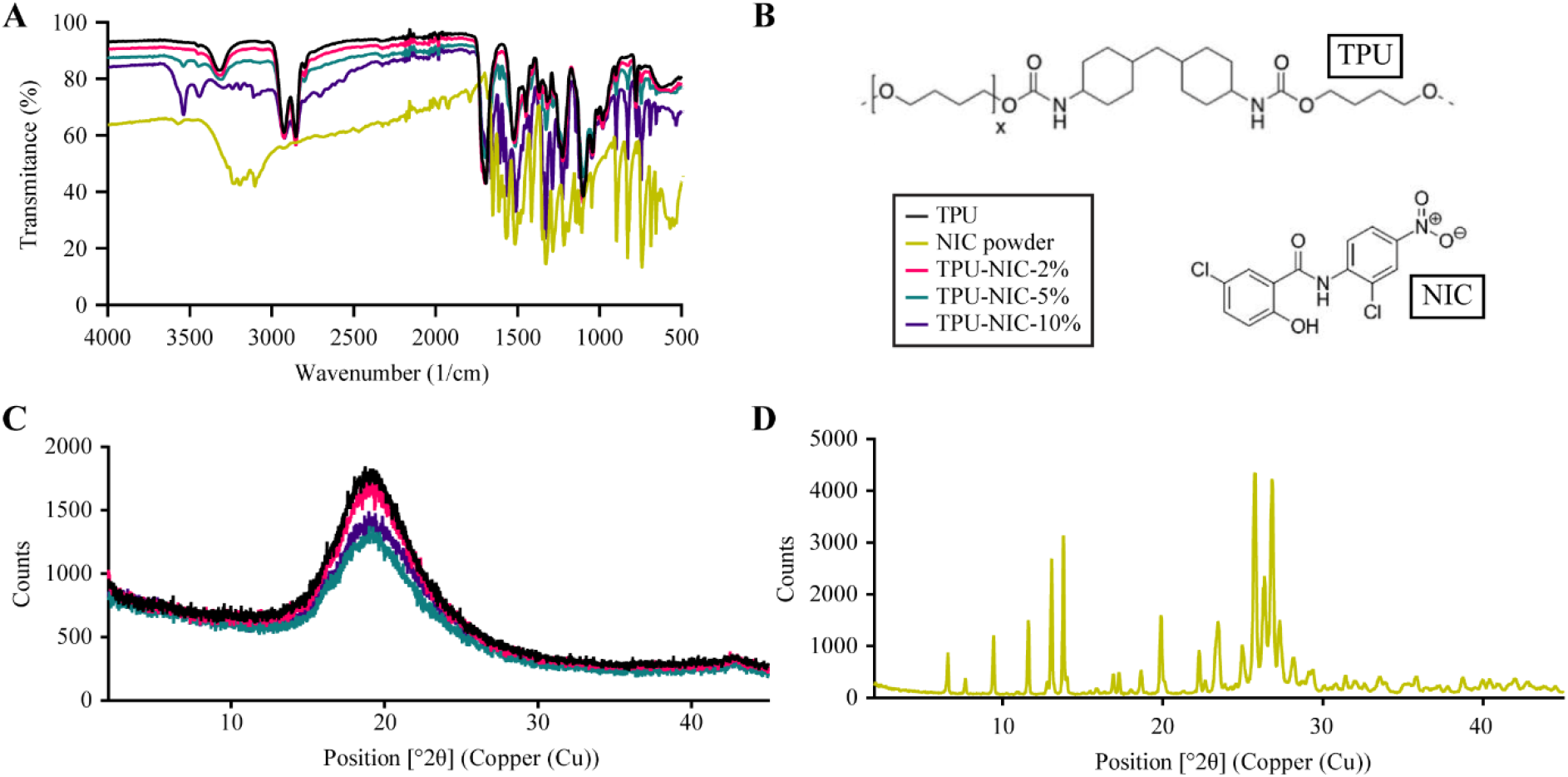
(A) FTIR spectra shown as the transmittance (%) of NIC, NIC-loaded and non-loaded TPU catheters (*n* = 3), (B) Molecular structure of TPU and NIC used in this study, (C) X ray powder diffraction pattern of non-loaded TPU catheters and TPU catheters loaded with 2, 5 and 10 % (w/w) NIC and (D) NIC plain drug.

The XRD patterns of the non-loaded and NIC-loaded TPU tubes and pure NIC powder are reported in the Figure 4C and D. NIC powder exhibited the characteristic peaks corresponding to the crystal form of the molecule [41]. On the contrary, the XRD pattern of TPU indicated its amorphous nature [42]. The TPU tubes loaded with NIC showed an XRD pattern without the sharp NIC diffraction peaks, which suggests a transformation of crystalline NIC to its amorphous form, this phase change was also reported by Jara *et al*. when hot-melt extruding NIC with the polymer poly(1-vinylpyrrolidone-co-vinyl acetate) (PVP–VA) at 180°C for the production of enteric formulations with increased bioavailability [43].

### 3.3. NIC thermostability

Heating of NIC at 180°C did not affect the antibacterial activity of the drug as judged from the MIC and MBC against MSSA and MRSA strains (*Table 1*). This indicates that NIC is thermally stable at the processing temperature and is expected not to lose its antimicrobial activity during extrusion at 180°C.

**Table 1:** MIC and MBC of non-heated NIC, and heated NIC at 180°C

| Bacteria | Niclosamide (µg/mL) |  |  |  |
| --- | --- | --- | --- | --- |
|  | Non heated |  | Heated (180°C) |  |
|  | MIC | MBC | MIC | MBC |
| MSSA | 0.0625 | 0.125 | 0.0625 | 0.125 |
| MRSA | 0.125 | 0.125 | 0.125 | 0.125 |

The estimated amount of NIC (μg) loaded per 1 cm catheter segment was 258 ± 3, 573 ± 5 and 1,110 ± 7 μg, for tubes loaded with 2, 5 and 10 % NIC, respectively. All NIC loaded catheter segments showed an initial burst release of NIC in the first 24 h, followed by a gradual and sustained release over 27 days (*Figure 5*A). Overall, the 2, 5 and 10 % NIC-loaded catheters showed similar drug release kinetics over time (*Figure 5*A-B). During the first 10 days, about 70 % of NIC was released from the catheter segments loaded with 10 % of NIC, while around 80 % of the drug was released from catheter segments loaded with 2 and 5 % of NIC. After 20 days, catheters loaded with 10 % of NIC released 80 % of the drug whereas a release of 90 % was achieved for catheters with 2 and 5 % of NIC (*Figure 5*A-B).

**Figure 5.**
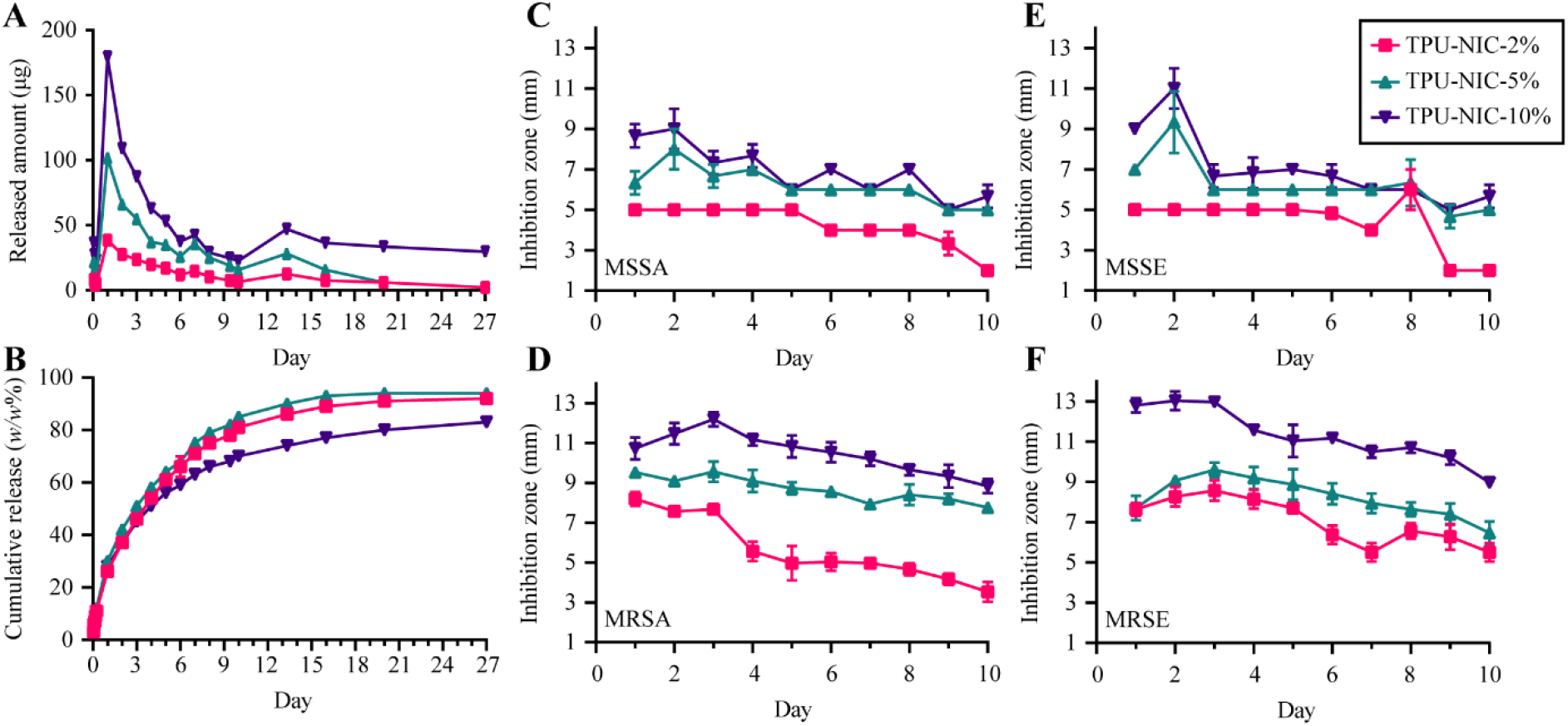
(A) Amount of NIC released (in μg) and (B) cumulative release (in *w/w*% of original NIC loading) over time up to 27 days, ZOI diameter (in mm) of (C) MSSA, (D) MRSA, I MSSE and (F) MRSE around NIC-loaded catheter segments transferred to fresh plates each day for 10 consecutive days (*n* = 3).

### 3.4. Antimicrobial activity of NIC over time

All catheter segments loaded with 2, 5 and 10 % NIC showed a detectable zone of inhibition (ZOI) of bacterial growth for MSSA, MRSA, MSSE and MRSE for 10 consecutive days (*Figure 5*C-F). As expected, non-loaded catheter segments did not produce any ZOI. For the first 5 days, catheters segments loaded with 2 % of NIC produced a ZOI between 4 and 8 mm depending on the bacterial strain. On subsequent days, the ZOI was gradually reduced reaching in some strains a minimum diameter of 2 mm at 10 days. Catheter segments loaded with 5 % of NIC exhibited a larger ZOI, ranging from 8 to 12 mm during the first 5 days. On the last day of experimentation, the ZOI produced for all the strains tested ranged from 6 to 10 mm. Compared to catheters with other loading percentage, catheter segments loaded with 10 % of NIC showed the largest inhibition zones after 10 days. These results showed that released NIC from the catheters exhibited *in vitro* antimicrobial activity overtime.

### 3.5. Evaluation of antimicrobial properties of TPU-catheters loaded with NIC

The *in vitro* antibacterial activity of NIC-loaded catheters was assessed using MSSA, MRSA, MSSE, MRSE strains (*Figure 6*A-H).

**Figure 6.**
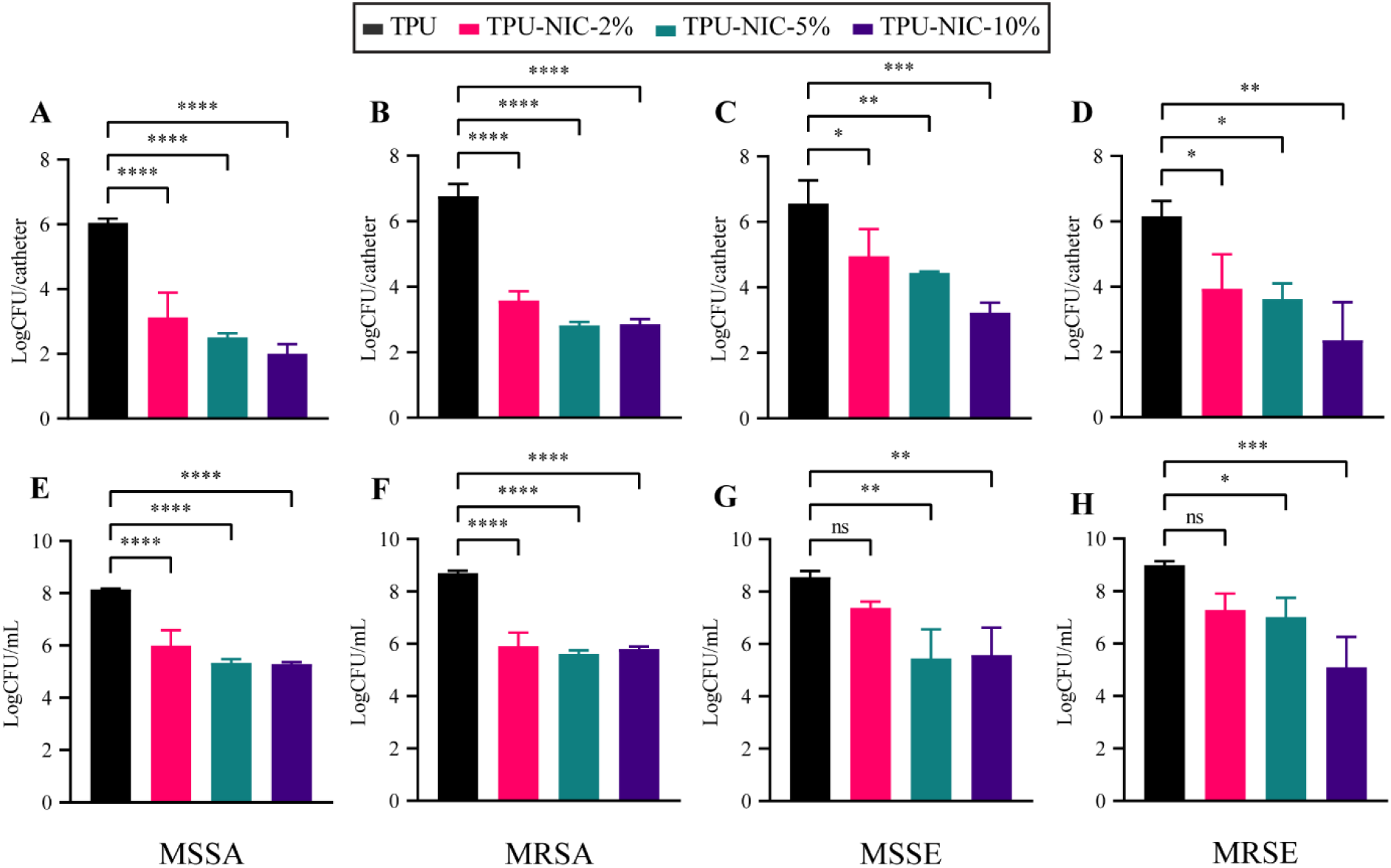
*In vitro* activity of NIC-loaded catheters against biofilm formation and planktonic growth in the surrounding medium after inoculation with 5×10^6^ CFU of different *S. aureus* and *S. epidermidis* strains. Top row: biofilm growth (24 h) on catheters segments and bottom row: Planktonic bacterial growth in surrounding media (24 h) of A, E) MSSA; B, F) MRSA; C, G) MSSE and D, H) MRSE. Each group was analyzed in triplicate and data expressed as mean LogCFU with SD. * Indicates a *p*-value of 0.01 to 0.05, ** indicates a *p*-value of 0.001 to 0.01, *** *p*-value of 0.0001 to 0.001, **** indicates a *p* value < 0.0001, ns indicates a non-significant *p*-value.

Biofilm formation was significantly reduced on the surface of all NIC-loaded catheter segments, with reductions higher than 2 LogCFU for both tested *S. aureus* strains (*Figure 6*A-B). Planktonic growth around all NIC-loaded catheters was reduced when compared against the non-loaded group (*Figure 6*E-F). However, similar reductions on the bacterial load were observed for all the drug loadings, indicating a possible bacteriostatic effect of released NIC as the LogCFU/catheter values are similar to those of the inoculum. The biofilm formation for both *S. epidermidis* strains on all NIC-loaded catheters segments was significantly reduced (> 1.6 LogCFU reductions) (*Figure 6*C-D) whereas corresponding planktonic bacterial growth was only reduced by catheter segments loaded with NIC at 5 and 10 % (*Figure 6*G-H). The reduction of the bacterial burden on catheter segments challenged with MSSA was confirmed by visualization with SEM (*Figure 7*A-C). In non-loaded catheter segments, a high number of bacterial cells were attached to the intraluminal surface forming a biofilm structure. In contrast, the catheter segments loaded with 2 and 5 % of NIC showed only few bacteria adhered to the surface.

**Figure 7.**
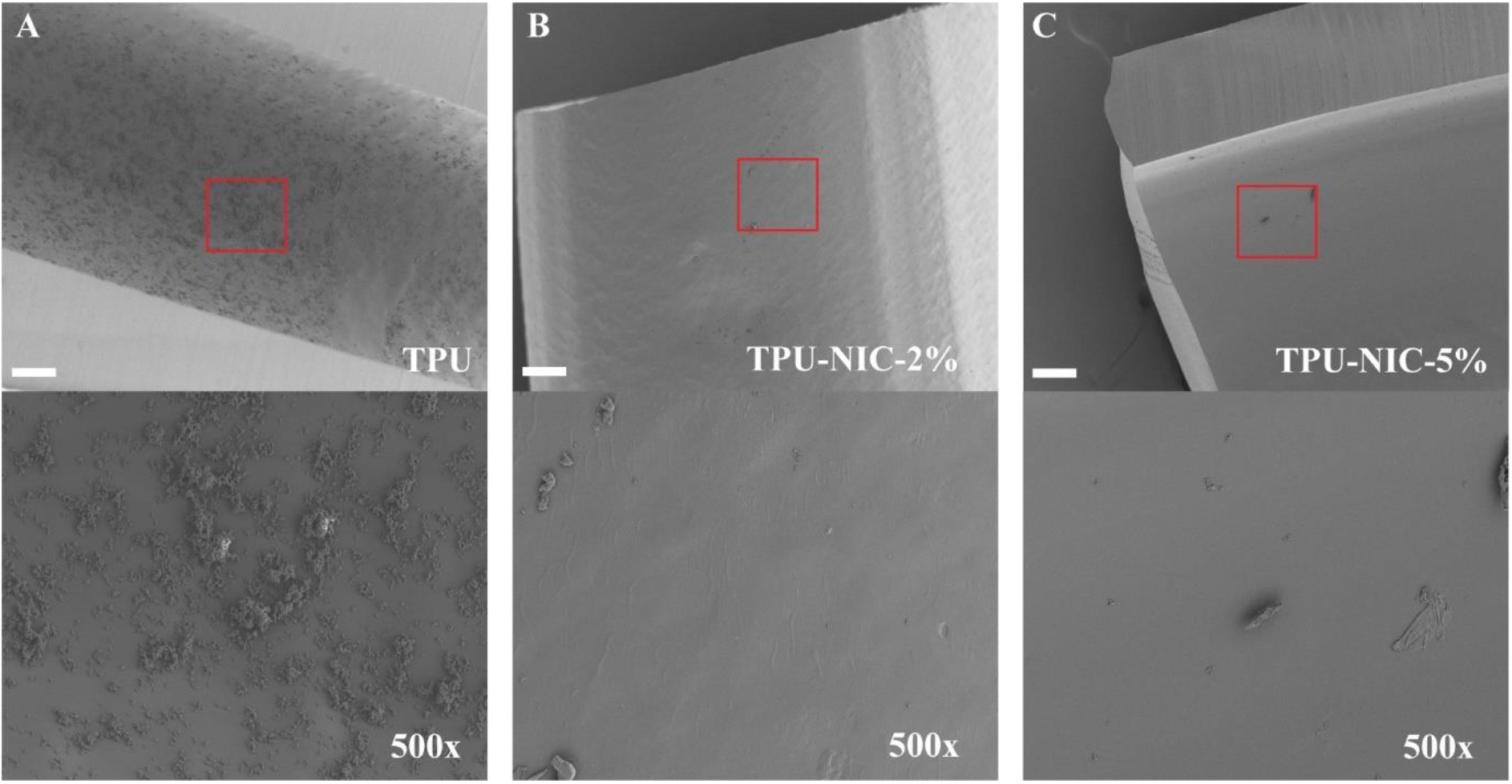
(A-C) Representative SEM images of 24 h biofilm formation of MSSA on catheter surfaces. Low magnification (top row, 100×) and higher magnification views (bottom row, 500×) of longitudinally sectioned catheter segments showing intraluminal bacterial colonization of TPU (A) and TPU loaded with 2 and 5 % of NIC (B and C).

### 3.6. *In vivo* evaluation of TPU-catheters loaded with NIC in a BAI model

As the 10 % NIC may affect the mechanical properties of TPU and to avoid potential toxic effects, *in vivo* efficacy of 2 and 5 % NIC loaded catheters was assessed in a BAI model against the MSSA strain (*Figure 8*A-B). On day 1, a significantly lower bacterial load was detected on both NIC loaded catheters (4.29 and 3.40 LogCFU/catheter for 2 and 5 % loadings, respectively) with respect to non-loaded catheter segments (6.06LogCFU/catheter) (p < 0.0001). Additionally, a significant reduction of 1.24 LogCFU in the bacterial counts was also observed in the surrounding tissue of 5 % NIC loaded segments (p < 0.01). On day 3 and 7 post infection, bacterial burden of both NIC-loaded catheters and in the respective surrounding tissues was progressively reduced over-time. Finally, the maximum effect was observed after 14 days when the bacterial load was greatly reduced (~5.7 Log CFU reduction p< 0.0001) on both NIC loaded catheters which was in line with further bacterial reduction of ~4.2 Log CFU (p < 0.0001) in corresponding surrounding tissue (*Figure 8*A-B).

**Figure 8.**
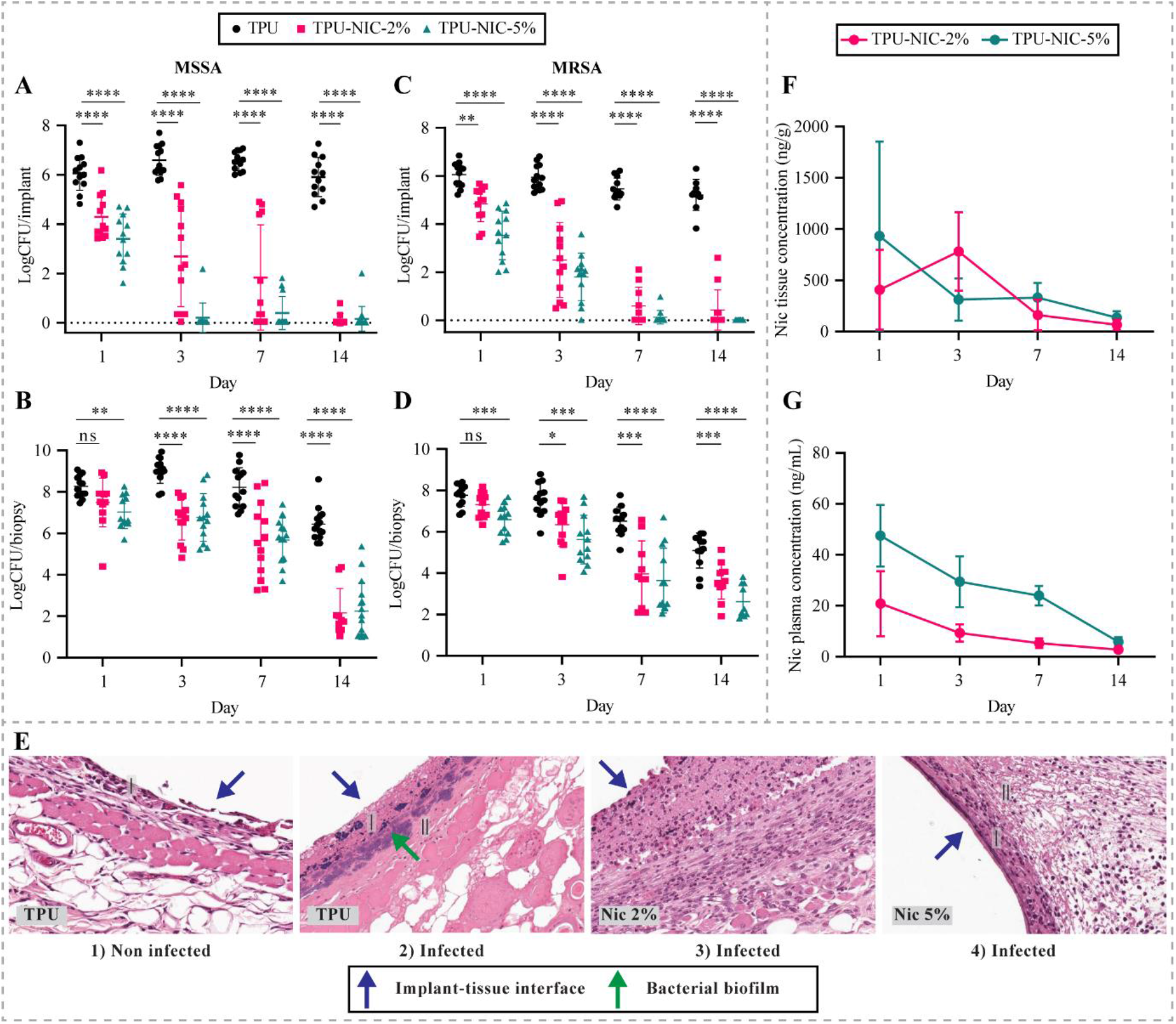
Antibacterial evaluation of TPU catheters loaded with 2 and 5 % of NIC at 1, 3, 7 and 14 days after implantation and infection in the mouse BAI model. *In vivo* efficacy of NIC-loaded catheters on MSSA colonization of A) catheters and B) peri-catheter tissue. Efficacy of NIC-loaded catheters on MRSA colonization of C) catheter and D) peri-catheter tissue. E) Histological analysis of MSSA subcutaneous biofilm infections in peri-catheter tissue sections of animals implanted with non-loaded and NIC-loaded catheters (I) Infiltration of neutrophiles/macrophages, (II) Fibrin deposition. Mice carrying non-loaded catheter segments without infection were used for baseline histological observations. Pharmacokinetic profiles of NIC released from catheters loaded with 2 and 5 % NIC over 14 days. Concentration of NIC F) in peri-catheter tissue and G) in plasma. * Indicates a *p*-value of 0.01 to 0.05, ** indicates a *p*-value of 0.001 to 0.01, *** *p*-value of 0.0001 to 0.001, **** indicates a *p* value < 0.0001, ns indicates a non-significant *p*-value.

In a subsequent experiment, mice implanted with catheter segments loaded with 2 and 5 % of NIC were challenged with the MRSA strain (*Figure 8*C-D). On day 1 after infection, the bacterial load on 2 and 5% NIC catheters (4.29 and 3.40 LogCFU/catheter; p < 0.001 and p < 0.0001, respectively) was significantly lower if compared to unloaded catheters (6.06 LogCFU/catheter). Moreover, bacterial counts in the surrounding tissue of 5 % NIC catheters was significantly reduced by a value of 1.23 LogCFU (p < 0.01), as similarly observed with the MSSA strain. On the following days, the bacterial burden on both NIC loaded segments and in the surrounding tissue gradually decreased over time until day 14. (*Figure 8*C-D). Few colonies of 2 and 5 % NIC groups, isolated at day 14, showed increased MIC values for NIC (2 colonies = MIC 1 μg/mL, 1 colony = MIC 2 μg/mL, and 1 colony = MIC 8 μg/mL). The MIC recorded for the original strain was replicated for most of the isolated colonies (i.e. 56 colonies MIC 0.25 μg/mL). Finally, no increase in terms of MIC values of NIC was observed in colonies isolated from non-loaded TPU groups.

Histology of tissue surrounding the catheters was performed on tissue samples from mice infected with MSSA and sacrificed on day 3 (*Figure 8*E). In mice implanted with non-infected non-loaded catheter segments without infection, the inflammation reaction was characterized by a small number of neutrophils and macrophages, with minimal fibrin deposition restricted to peri-catheter tissue. In addition, no necrosis was observed in this group (*Figure 8*E-1). On the other hand, in biopsies from mice with infected non-loaded catheter segments, the inflammation was characterized by the presence of abundant neutrophils and less macrophages which were embedded in abundant edema and fibrin. Extensive necrosis was observed around the catheter segments, variably extending into the surrounding tissues. Moreover, a full-thickness coagulative skin necrosis was observed in this group (*Figure 8*E-2). Intralesional bacterial biofilm formation was observed within the necrotic debris that surrounded the catheter segment. Additionally, features such as epidermal hyperplasia, furunculosis and skin ulceration were registered in the tissue. Animals with NIC-loaded catheter segments showed morphological changes similar to those present in the infected mice with non-loaded catheters, however the severity was not as marked. Additionally in the same biopsies, we observed a moderate number of neutrophils and macrophages with moderate fibrin deposition. Furthermore, a minimal number of necrotic debris was found in the surrounding tissue and no bacterial aggregates were observed, which agrees with the observed quantitative decrease in the bacterial load in surrounding tissue.

The NIC distribution *in situ* in the surrounding tissue is shown in Figure 8F. An initial NIC level of 407 ± 389 and 932 ng/g ± 922 of tissue was recorded in mice bearing catheter segments with 2 and 5 % NIC, respectively. After 3 days, NIC levels of 781 ± 382 ng/g and 312 ± 206 ng/g (no significant difference between both groups) were recorded in the surrounding tissue of catheter segments loaded with 2 and 5 % NIC, respectively. On the following experimental days, the *in situ* NIC concentration decreased in the surrounding tissue, reaching the lowest values of 65 ± 51 ng/g (NIC 2 %) and 135 ± 62 ng/g (NIC 5 %) on day 14. The *in vivo* level of NIC in plasma is shown in Figure 8G. After 1 day, NIC plasma levels of 20 and 47 ng/mL were measured for catheters with 2 and 5 % of NIC, respectively. On the consecutive time points, the NIC plasma levels for both NIC loadings gradually dropped over time, reaching levels at the limit of detection on day 14.

## 4. Discussion and conclusion

Catheters are essential medical devices which are mainly used for diagnosis, prevention, and treatment of many diseases and health conditions [1]. Catheters are mainly fabricated by HME process in which medical grade polymers are fed into an extrusion machine, then heated up to melting and finally extruded through a nozzle [44]. Anti-infective catheters can be produced by loading the active pharmaceutical ingredient (API) within the polymeric matrix of the catheter. This can be done during the HME processes, however, the thermal stability of the loaded API should be taken into consideration to avoid any possible degradation [45].

We developed a novel NIC eluting catheter adapting the methodology previously described by Shaqour *et al*. [29] to prevent ventilator-associated pneumonia. We used NIC, as a repurposed antibacterial drug, to minimize the potential development of antibiotic resistance which currently is a rising global health issue [46]. Moreover, the potential antibacterial activity of NIC has been tested against multi-drug resistant bacteria such as methicillin-resistant *S. aureus* and *S. epidermidis*, two of the most prominent pathogens found in medical device-related infections [13, 14]. Furthermore, NIC has not only been proven biologically active in different preclinical models but also not toxic in clinical trials [15, 21, 47–50].

We confirmed that NIC can withstand processing temperatures (up to 180°C) without undergoing degradation [43] and any detrimental changes in its *in vitro* antibacterial activity. Moreover, NIC has a well-studied safety profile thereby it has been tested to treat different medical indications using preclinical models [24, 51–56]. Considered together, these results prove NIC’s suitability to fabricate anti-infective medical devices by using not only HME but also different processes involving temperatures at least up to 180°C such as melt extrusion 3D-printing technologies. After proving the suitability of NIC for HME applications, NIC-loaded TPU catheters were fabricated at loadings of 2, 5 and 10 % NIC (*w/w*). Overall, the addition of NIC at 2 and 5 % (*w/w*) did not alter the mechanical properties of the TPU fibers which would likely be the same for NIC-loaded catheters. The incorporation of NIC into the polymeric matrix of TPU catheters was confirmed by the appearance of new functional groups in the FTIR spectra. These changes in NIC-loaded TPU likely are due to covalent modifications, while the shift of spectral bands characteristic of the TPU or the immobilized NIC may indicate non-covalent interaction [57, 58]. Moreover, NIC underwent a phase transition from crystalline to an amorphous form within the NIC-loaded catheters. This drug phase transition is likely caused by the combined effect of solvent casting and the HME process during catheter production [43, 59]. The amorphous form of NIC improves the drug’s bioavailability, which in turn is pivotal for a successful delivery of the drug within a host and to exert the intended antibacterial effect [59].

Catheters loaded with NIC presented a steady drug release over 27 days. In similar drug-loaded implants, the drug release is mainly mediated by passive diffusion, which governs how the drug migrates towards the external media [60]. In drug delivery systems governed by diffusion, the release kinetics relies on the solubility of the drug in the polymer, the diffusion coefficient of the drug, the drug loading, and the degradation of the polymer[61]. In fact, we observed that the total amount of released NIC to the media was dependent on the drug loading.

The *in vitro* antibacterial properties of NIC-loaded catheter segments were demonstrated by both a disc diffusion assay and the quantification of biofilm-forming bacteria on catheter surfaces and planktonic bacterial growth in liquid media [62]. Not only did the NIC prevent biofilm formation, the apparent release of NIC in the medium inhibited the bacteria to grow. The released concentrations of NIC however did not kill the planktonic bacteria, since their numbers did not significantly decrease relative to the inoculum, suggesting a bacteriostatic activity of released NIC.

Based on the *in vitro* studies, we next investigated whether the protective activity of NIC-loaded catheters with 2 and 5 % of NIC would translate to *in vivo* efficacy using a BAI model. In prior studies, this *in vivo* model has been proven useful to fast track the development of new antimicrobials and anti-infective medical devices to combat medical device-related infections [63]. In this *in vivo* model, the NIC-loaded catheters exhibited a similar antibacterial efficacy against both MSSA and MRSA strains for at least 14 days. The *in vivo* efficacy was reflected in significantly reduced bacterial colonization of both the catheter segments and the peri-catheter tissue.

From the *in vivo* experiments with MRSA, a limited number of colonies re-cultured from the tissue of mice with 2 % or 5 % NIC-loaded catheter segments displayed elevated MIC values. As the antibacterial indication is not approved for NIC, there are no data available of resistance concentration breakpoints for clinical susceptibility. The shifts in the MIC of NIC could be due to either emergence of resistance or tolerance, and in case of the later, it could be circumvented by delivering therapeutically attainable NIC concentrations to the infection site. Apparently, the elevated MIC had not resulted in dominance of this trait in the bacteria in the tissue since most of the retrieved colonies had lower MIC values. Resistance development of Gram-positive bacteria to NIC has to the best of our knowledge not been reported until now but should be kept in mind as a possibility of prolonged *in vivo* exposure to concentrations released, which are decreasing over time. Therefore, we strongly suggest performing NIC susceptibility assays with bacteria re-cultured from *in vivo* models with NIC as the antimicrobial agent. On the other hand, Gram-negative bacteria possess inherent resistance to NIC, mediated by efflux pumps and nitro-reduction [64]. Interestingly, it has been reported that co-administration of NIC and colistin exerted *in vivo* synergistic antibacterial efficacy against resistant *Pseudomonas aeruginosa* in a skin abscess infection model, which is a difficult to treat infection with standard antibiotics alone [64]. Therefore, the development of catheters containing NIC and synergistic antibiotics holds potential to decrease the likelihood of emergence of newly NIC tolerance/resistance while increasing the antibacterial efficacy against Gram-positive and Gram-negative bacteria.

The reduction of bacterial colonization in surrounding tissue was also confirmed by histological analysis, which is relevant since peri-catheter tissue is a niche allowing infecting bacteria to survive in medical device-related infections [65, 66]. Also, NIC-loaded catheters hold the potential to elicit a complete anti-infective protection against bacteria coming from the skin or emanating from contaminated hubs [5, 67, 68]. In addition, our data suggest that NIC is rapidly released and retained into peri-catheter tissue, since the local tissue concentration was several folds higher than in plasma. Importantly these NIC plasma levels have been reported to not exert systemic toxicity [21, 47, 48]. The higher NIC levels in tissue indicate that NIC likely slowly diffuses through the tissue and into the blood vessels. From similar reported *in vivo* release kinetics, we can hypothesize that once in blood NIC undergoes a rapid systemic clearance which in fact will minimize the systemic concentrations of NIC [69]. In addition, considering the amount of NIC in the loadings of 2 and 5 % per 1 cm catheter, and the initial mean weight of the mice (~20 g) at the beginning of the experiments, we approximately administered to the mice a NIC dose of 12.5 mg/kg and 31.25 mg/kg per a NIC loading of 2 and 5 % respectively. Different preclinical studies have reported that daily subcutaneous NIC doses of 1 and 10 mg/kg and intraperitoneal NIC doses of 10, 20, 30 and 50 mg/kg did not exert any toxic effect in mice after several days of treatment [64, 70–74]. Moreover, the acute oral toxicity LD50 value for mice is 1,500 mg/kg while the subcutaneous LD50 value is >1,000 mg/kg[75]. In its anthelmintic indication NIC is administered orally at a dose of 2,000 mg/patient and is well tolerated by human patients [76]. Furthermore, the amount of NIC loaded in the catheters is gradually released into the peri-catheter tissue which modulates the real concentration of bioavailable NIC overtime and therefore minimizes the exposure to potential systemic toxic NIC levels.

Although the BAI model does not completely mimic a catheterization in mice, this model offers relevant valid *in vivo* information on the functionality of NIC-loaded catheters to resist *S. aureus* infection in potential clinical conditions [63]. Moreover, our results suggest a potential clinical application for NIC-loaded TPU catheters, as NIC was able to combat infection by the MRSA strain tested, and MRSA strains are known to cause most of the life-threatening staphylococcal infections [53, 54]. Furthermore, the approach followed in this research work could be applied for developing novel NIC-functionalized medical devices to prevent or treat staphylococcal infections such as those associated with wounds, heart valves, urinary catheters, intra-uterine devices, voice prostheses, endotracheal tubes, and prosthetic implants. In future studies it would be interesting to use combinations of NIC and other antibacterial agents to generate broad-spectrum antibacterial catheters. In conclusion, this research work serves as a proof of concept of the potential use of NIC to develop medical devices using high-temperature manufacturing technologies, for preventing medical device-related infections.

## 5. Acknowledgment

This research was funded by the research project PRINT-AID, the EU Framework Programme for Research and Innovation within Horizon 2020—Marie Skłodowska-Curie Innovative Training Networks under grant agreement No. 722467.

The authors would like to thank Prof. Christophe Vande Velde from the Intelligence in Processes, Advanced Catalysts and Solvents (iPRACS) research group, Antwerp University for allowing researchers to use the thermogravimetric analysis machine in his lab. In addition, Ana Criado and Rosella Defazio for their technical assistance in the histological experimentation (Evotec, Verona, Italy); Chiara Pignaffo and Stefano Fontana (Evotec, Verona Italy) for their contribution in the *in-vivo* release experiments. Similarly, Jhon Quintana and Giulio Giommarelli for their assistance with the *in vivo* experimentation. Additionally, Mr. Jean-Pierre Smet from the Material Science department for allowing researchers to use the tensile machine in his lab. We would also like to thank Edwin Scholl and Dr. Nicole van der Wel (Electron Microscopy Center Amsterdam (EMCA), Amsterdam UMC) for their technical assistance in the collection of the SEM images. We would also like to thank Dr. Bartłomiej Wysocki and Agnieszka Chmielewska for producing the coaxial nozzle that were used to extrude the catheters.

## 6. Author Contributions

Conceptualization A.V., B.S., C.G.-P., A.F., L.F., M.R., P.C., S.A.J.Z., B.V, K.B, V.J.A.C. and W.S.; investigation A.V., B.S., C.G.-P and E.C.; writing—original draft preparation A.V., B.S. and C.G.-P.; writing—review and editing A.F., L.F., V.J.A.C., M.R., P.C., S.A.J.Z., E.C. and P.C.; supervision A.F., L.F., P.C., M.R., S.A.J.Z., B.V., K.B., V.J.A.C. and W.S.; funding acquisition A.F., L.F., B.V., K.B., P.C., S.A.J.Z., W.S. and P.C. All authors have read and agreed to the published version of the manuscript.

---

**Supplementary Table 1.** Instrumental parameters. The XRD analyses were run in transmission mode on an X-Ray Diffractometer equipped with an X’Celerator detector using a standard XRD method.

| Instrumental Parameter | Value |
| --- | --- |
| 2-theta range | 2-45° |
| Step size [°2-theta] | 0.0167 |
| Time per step [sec] | 59.690 sec |
| Scan Mode | Continuous |
| Sample Movement | Spinning, 1.0sec rotation time |
| Wavelength [nm] | Cu $K\alpha_1=1.54060$ $K\alpha_2=1.54443$ |
| X-ray Mirror | Inc. Beam Cu W/Si focusing MPD, Acceptance Angle 0.8°C, Length 55.3 mm |
| Incident Divergence Slit | Slit Fixed 1/2° |
| Incident Antiscatter Slit | Slit Fixed 1/2° |
| Incident Beam Mask | 10 mm |
| Beam Stop | Beam Stop for Transmission Spinner |
| Temperature/RH | Room temperature |
| Soller Slits | 0.02 rad on Incident and Diffracted beam |
| Detector type | X'Celerator (active length 2.122°) |
| Sample holder | Transmission sample holder. Samples are mounted between Mylar film |
| Configuration | Transmission |
| Generator voltage/current | 40 KV / 40 mA |

**Supplementary figure 1.**
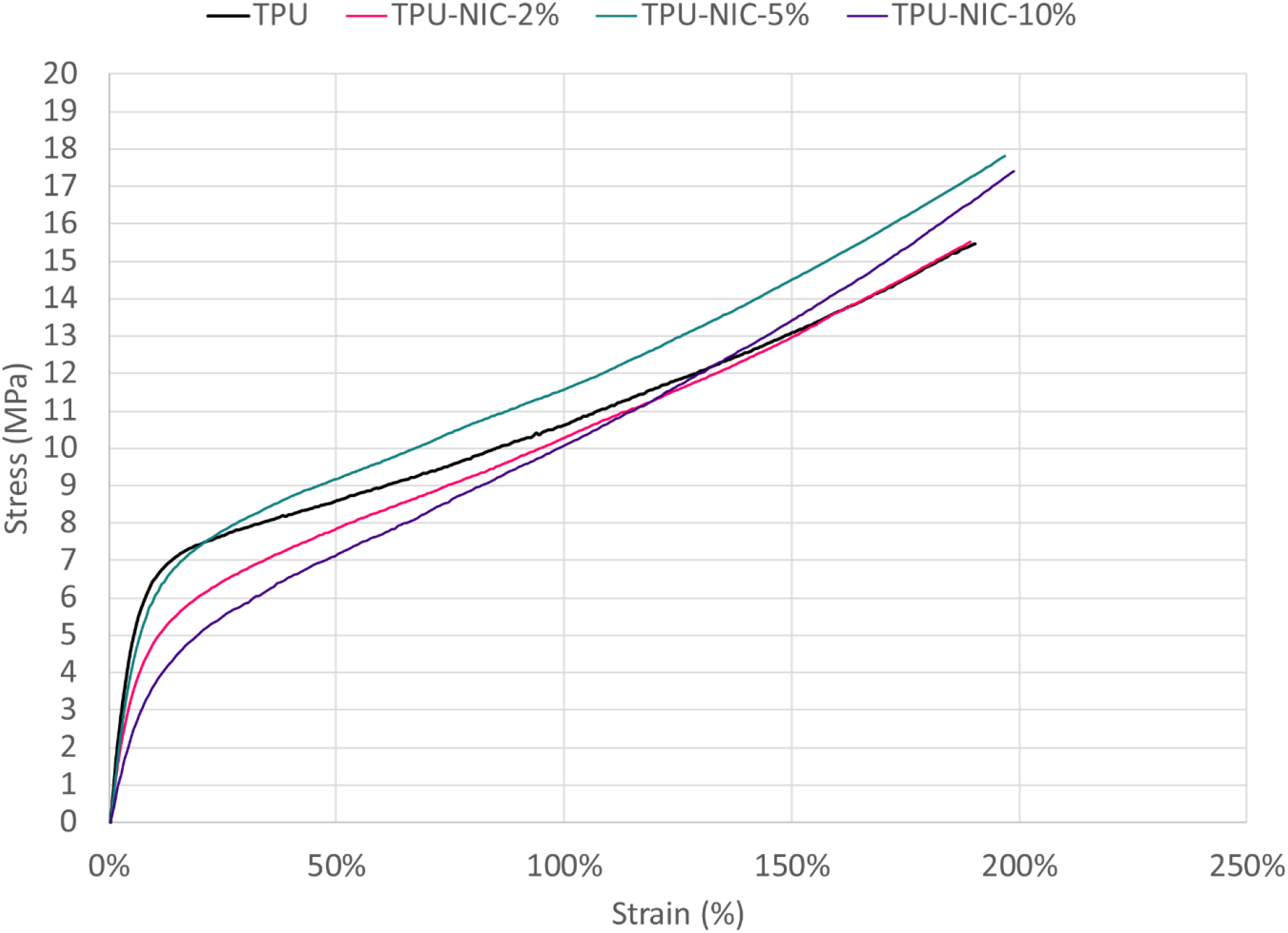
Representative stress strain curve generated from tensile test.

**Supplementary figure 2.**
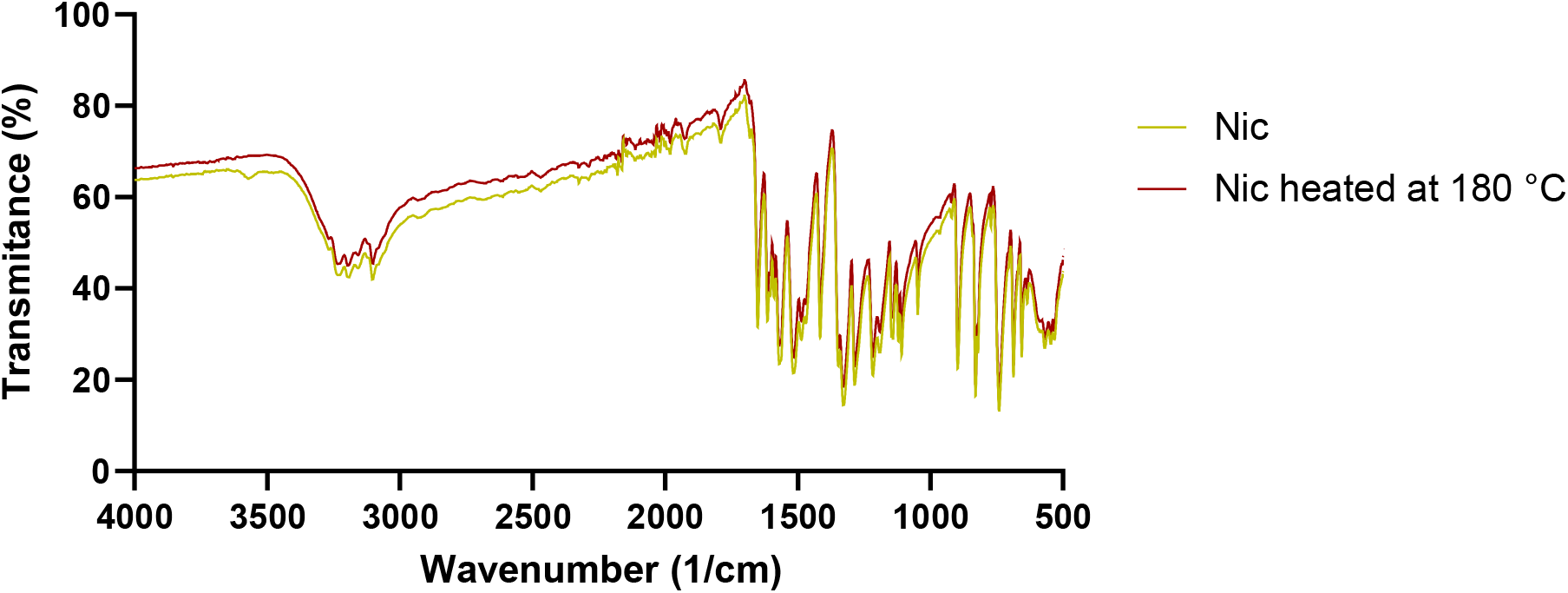
FTIR spectra of neat Nic (yellow line) and heated for 5 min at 180 °C (red line).

